# Wolf-Hirschhorn Syndrome-associated genes are enriched in motile neural crest and affect craniofacial development in *Xenopus laevis*

**DOI:** 10.1101/471672

**Authors:** Alexandra Mills, Elizabeth Bearce, Rachael Cella, Seung Woo Kim, Megan Selig, Sangmook Lee, Laura Anne Lowery

**Affiliations:** Boston College, Biology Depart, Chestnut Hill, MA, USA

**Keywords:** craniofacial development, developmental disorders, 4p-, Wolf-Hirschhorn Syndrome, WHSC1, WHSC2, LETM1, TACC3, neural crest, disease model

## Abstract

Wolf-Hirschhorn Syndrome (WHS) is a human developmental disorder arising from a hemizygous perturbation, typically a microdeletion, on the short arm of chromosome four. In addition to pronounced intellectual disability, seizures, and delayed growth, WHS presents with a characteristic facial dysmorphism and varying prevalence of microcephaly, micrognathia, cartilage malformation in the ear and nose, and facial asymmetries. These affected craniofacial tissues all derive from a shared embryonic precursor, the cranial neural crest, inviting the hypothesis that one or more WHS-affected genes may be critical regulators of neural crest development or migration. To explore this, we characterized expression of multiple genes within or immediately proximal to defined WHS critical regions, across the span of craniofacial development in the vertebrate model system *Xenopus laevis*. This subset of genes, *WHSC1, WHSC2, LETM1*, and *TACC3*, are diverse in their currently-elucidated cellular functions; yet we find that their expression demonstrates shared tissue-specific enrichment within the anterior neural tube, pharyngeal arches, and later craniofacial structures. We examine the ramifications of this by characterizing craniofacial development and neural crest migration following individual gene depletion. We observe that several WHS-associated genes significantly impact facial patterning, cartilage formation, pharyngeal arch migration, and neural crest motility, and can separately contribute to forebrain scaling. Thus, we have determined that numerous genes within and surrounding the defined WHS critical regions potently impact craniofacial patterning, suggesting their role in WHS presentation may stem from essential functions during neural crest-derived tissue formation.

**Author Summary:** Wolf-Hirschhorn Syndrome (WHS), a developmental disorder caused by small deletions on chromosome four, manifests with pronounced and characteristic facial malformation. While genetic profiling and case studies provide insights into how broader regions of the genome affect the syndrome’s severity, we lack a key component of understanding its pathology; a basic knowledge of how individual WHS-affected genes function during development. Importantly, many tissues affected by WHS derive from shared embryonic origin, the cranial neural crest. This led us to hypothesize that genes deleted in WHS may hold especially critical roles in this tissue. To this end, we investigated the roles of four WHS-associated genes during neural crest cell migration and facial patterning. We show that during normal development, expression of these genes is enriched in migratory neural crest and craniofacial structures. Subsequently, we examine their functional roles during facial patterning, cartilage formation, and forebrain development, and find that their depletion recapitulates features of WHS craniofacial malformation. Additionally, two of these genes directly affect neural crest cell migration rate. We report that depletion of WHS-associated genes is a potent effector of neural crest-derived tissues, and suggest that this explains why WHS clinical presentation shares so many characteristics with classic neurochristopathies.

## Introduction

Wolf-Hirschhorn Syndrome (WHS) is a developmental disorder characterized by intellectual disability, delayed pre- and post-natal growth, heart and skeletal defects, and seizures [1–4]. A common clinical marker of WHS is the “Greek Warrior Helmet” appearance; a facial pattern with a characteristic wide and flattened nasal bridge, a high forehead, drastic eyebrow arches and pronounced brow bones, widely spaced eyes (hypertelorism), a short philtrum, and an undersized jaw (micrognathia). The majority of children with the disorder are microcephalic, and have abnormally positioned ears with underdeveloped cartilage. Comorbid midline deficits can occur, including cleft palate and facial asymmetries [1]. Craniofacial malformations make up one of the most prevalent forms of congenital defects [5,6], and can significantly complicate palliative care and quality of life [7]. Given the commanding role of cranial neural crest (CNC) cells in virtually all facets of craniofacial patterning, craniofacial abnormalities are typically attributable to aberrant CNC development [6,8]. A striking commonality in the tissues that are impacted by WHS is that a significant number derive from the CNC. Despite this, little is known about how the vast diversity of genetic disruptions that underlie WHS pathology can contribute to craniofacial malformation, and no study has sought to characterize impacts of these genotypes explicitly on CNC behavior.

WHS is typically caused by small, heterozygous deletions on the short-arm of chromosome 4 (4p16.3), which can vary widely in position and length. Initially, deletion of a very small critical region, only partial segments of two genes, was thought to be sufficient for full syndromic presentation [9–13]. These first putative associated genes were appropriately denoted as Wolf-Hirschhorn Syndrome Candidates 1 and 2 (*WHSC1, WHSC2*) [9,11, 14-16]. However, children with WHS largely demonstrate 4p disruptions that impact not only this intergenic region between *WHSC1* and *WHSC2*, but instead affect multiple genes both telomeric and centromeric from this locus [17]. Focus was drawn to these broader impacted regions when cases were identified that neglected this first critical region entirely but still showed either full or partial WHS presentation, prompting the expansion of the originally defined critical region to include a more telomeric segment of WHSC1, and a new candidate, *LETM1* [4, 18]. These discrepancies are increasingly rectified by mounting evidence that true cases of the syndrome are multigenic [1,19-20]. Our emerging understanding of WHS as a multigenic developmental disorder necessitates its study as such—with a renewed focus on how the depletion of these genes combinatorially contribute to a collaborative phenotype. However, a central problem arises that entirely precludes this effort: we largely lack a fundamental understanding of how singular WHS-affected genes function in basic developmental processes. Furthermore, animal models of WHS-associated gene depletion have occurred across numerous species and strains, with no unifying model to offer a comparative platform. Given the disorder’s consistent and extensive craniofacial malformations, it seems especially prudent to establish whether these genes serve critical functions explicitly during processes governing craniofacial morphogenesis.

To this aim, we sought to perform a characterization of the contributions of four commonly WHS-affected genes, *WHSC1, WHSC2, LETM1*, and *TACC3*(1), during early craniofacial patterning in Xenopus laevis. We first examined expression profiles of these transcripts across early embryonic development, and notably, observed enrichment of all four transcripts in motile CNCs of the pharyngeal arches, which invites the hypothesis that they may impact neural crest development and migration. Knockdown (KD) strategies were then utilized to examine WHS-associated gene contributions to facial morphogenesis and cartilage development. We find that all KDs could variably affect facial morphology. Perhaps most notably, WHSC1 depletion increased facial width along the axis of the tragion (across the eyes or temples), recapitulating one feature of WHS craniofacial malformation. We performed both *in vivo* and *in vitro* CNC migration assays that illustrate that two of these genes (TACC3, WHSC1) can directly affect migrating pharyngeal arch morphology and CNC motility rates. Separately, as most of the examined transcripts also demonstrated enrichment in the anterior neural tube, we examined their impacts on embryonic forebrain scaling. We found that depletion of three of the four genes could additionally impact forebrain size. Together, our results support a hypothesis that WHS produces consistent craniofacial phenotypes (despite a vast diversity in genetic perturbations), in part due to numerous genes within the affected 4p locus performing critical and potentially combinatorial roles in neural crest migration, craniofacial patterning, cartilaginous tissue formation, and brain development. Furthermore, this work is the first to perform depletion of multiple WHS-affected genes on a shared, directly-comparable, laying an essential foundation for future efforts to model, integrate, or predict interactions of diverse genetic disruptions within the context of a multigenic syndrome.

## Results

### Numerous WHS-affected genes demonstrate enriched expression in the pharyngeal arches, early nervous system, and embryonic craniofacial structures

Pronounced and characteristic craniofacial dysmorphism is one of the most recognizable features of WHS-affiliated 4p16.3 microdeletions. Children with the disorder demonstrate a low-profile nasal bridge and prevalent lower forehead, with wide-set eyes and a short philtrum (together commonly referred to as the Greek Warrior’s Helmet presentation). Microcephaly and micrognathia are present with varying severity, and comorbidities commonly include facial asymmetries and cleft palate [21]. Given the commanding role of cranial neural crest (CNC) cell proliferation, migration, and differentiation in properly coordinated facial patterning of nearly all of these affected tissues, we hypothesized that certain WHS-affected genes could play critical roles in neural crest maintenance, motility, or specification, and that their depletion would thus disproportionately impact tissues derived from the neural crest.

We first performed coordinated examinations of spatiotemporal expression of commonly affected genes in the 4p16.3 locus across craniofacial development(2). To this end, we performed *in situ* hybridization with DIG-labeled antisense RNA probes against four genes within and proximal to the last defined WHS critical region (*WHSC1, WHSC2, LETM1*, and *TACC3*) (1). During early craniofacial morphogenesis at stage 25, we note enriched expression of *WHSC1, WHSC2*, and *TACC3* in the migrating pharyngeal arches (Fig. 2B, C, E), as their enrichment closely resembles the expression pattern of the CNC-enriched transcription factor Twist (Fig. 2A, F). Comparatively, *LETM1* (Fig. 2D) demonstrates ubiquitous expression. Interestingly, these transcripts are not significantly enriched in specified, premigratory neural crest (st. 16), with the exception of *TACC3* (Fig. S1). By stage 35, all four transcripts are enriched in pharyngeal arches (Fig. 2G-J); *LETM1* expression appears to reduce in neighboring tissues, while remaining selectively enriched in later stages of pharyngeal arch migration (Fig. 2I). There is also significant transcription of all four genes within the anterior neural tube. Later in tailbud stages, we note that some transcripts maintain enriched expression in the forebrain, most notably *WHSC2, while WHSC1, WHSC2*, and *LETM1* illustrate enrichment in tissues within the head and face (Fig. S1, EF, KL, QR, WX). Additionally, *WHSC1* and *LETM1* expression show potential overlap with cardiac tissue (Fig. S1E, Q).)

**Fig. 1.**
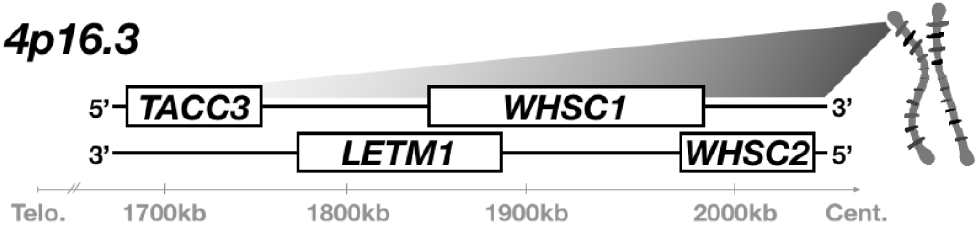
WHS is typically caused by heterozygous microdeletion of numerous genes within 4p16.3. A segment of this region is illustrated here. A microdeletion that spans at least *WHSC1*, *WHSC2*, and LETM1 is currently assumed to be necessary for full WHS diagnostic presentation; children affected by the disorder often possess larger deletions that extend further telomeric and impact additional genes, such as *TACC3*.

**Fig. 2.**
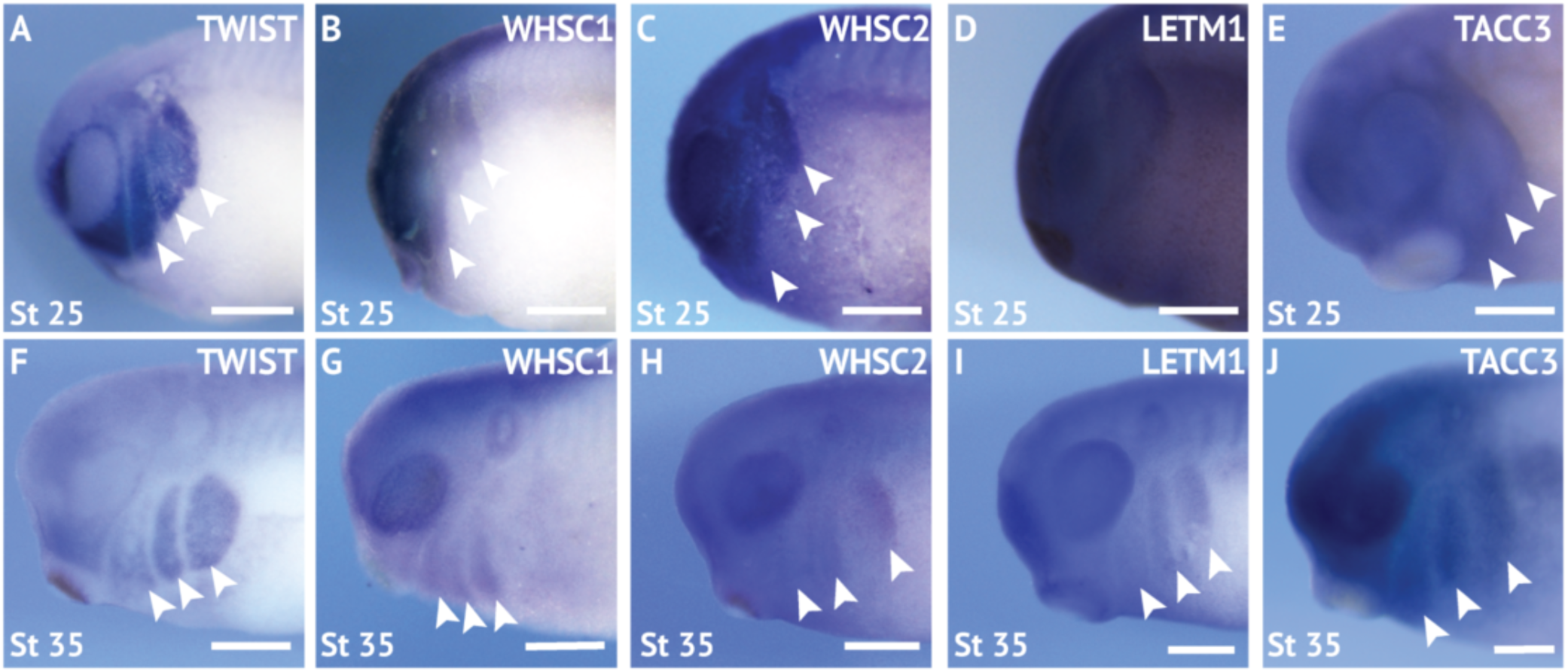
WHS related genes are expressed in the migrating neural crest cells during embryonic development. (A, F) Lateral views of whole mount *in situ* hybridizations for Twist, a CNC-enriched transcription factor. Arrows indicate the pharyngeal arches (PA). (B-E, G-J) *In situ* hybridizations for WHSC1, WHSC2, LETM1, and TACC3 demonstrate enrichment in motile PAs. Scalebar is 250μm.

### WHS-affected genes are critical for normal craniofacial morphology

Given that all four genes showed enrichment in migratory neural crest by stage 35, and most demonstrated enduring transcription in later craniofacial tissues, we hypothesized that their reduction may drive changes in craniofacial morphogenesis. To this end, we performed partial genetic depletions of all four genes individually, and performed morphometric analyses of craniofacial landmarks between WHS-associated gene depleted embryos and controls from the same clutch. Measurements to quantify facial width, height, midface area, and midface angle were performed as previously described [22] at stage 40.

Individual depletion of the examined WHS-affected genes demonstrated pronounced impacts on facial patterning (3). WHSC1 KD significantly increased facial width (Fig. 3F), and this increase accompanied a significant increase in facial area (Fig. 3H). WHSC1, LETM1, and TACC3 depletion conversely narrowed facial width at this axis (Fig. 3F), and additionally decreased facial area. None of these changes were proportional to facial height, which was unaffected by gene depletion. In nearly all cases, the distribution of facial features was normal. Only TACC3 depletion modestly affected the mid-face angle, a parameter describing the relationship between the eyes and mouth (Fig. 3I). Importantly, all facial phenotypes could be rescued by co-injection with full-length mRNA transcripts of their targets (Fig. S2), indicating that phenotypes were specific to WHS-associated gene depletion. Taken together, these results are consistent with a possibility that WHSC1 depletion may be sufficient to drive frontonasal dysmorphism, while WHSC2, LETM1, and TACC3 depletions may contribute to complex or epistatic interactions that mediate additional characteristic facial features of the developmental disorder.

**Fig. 3.**
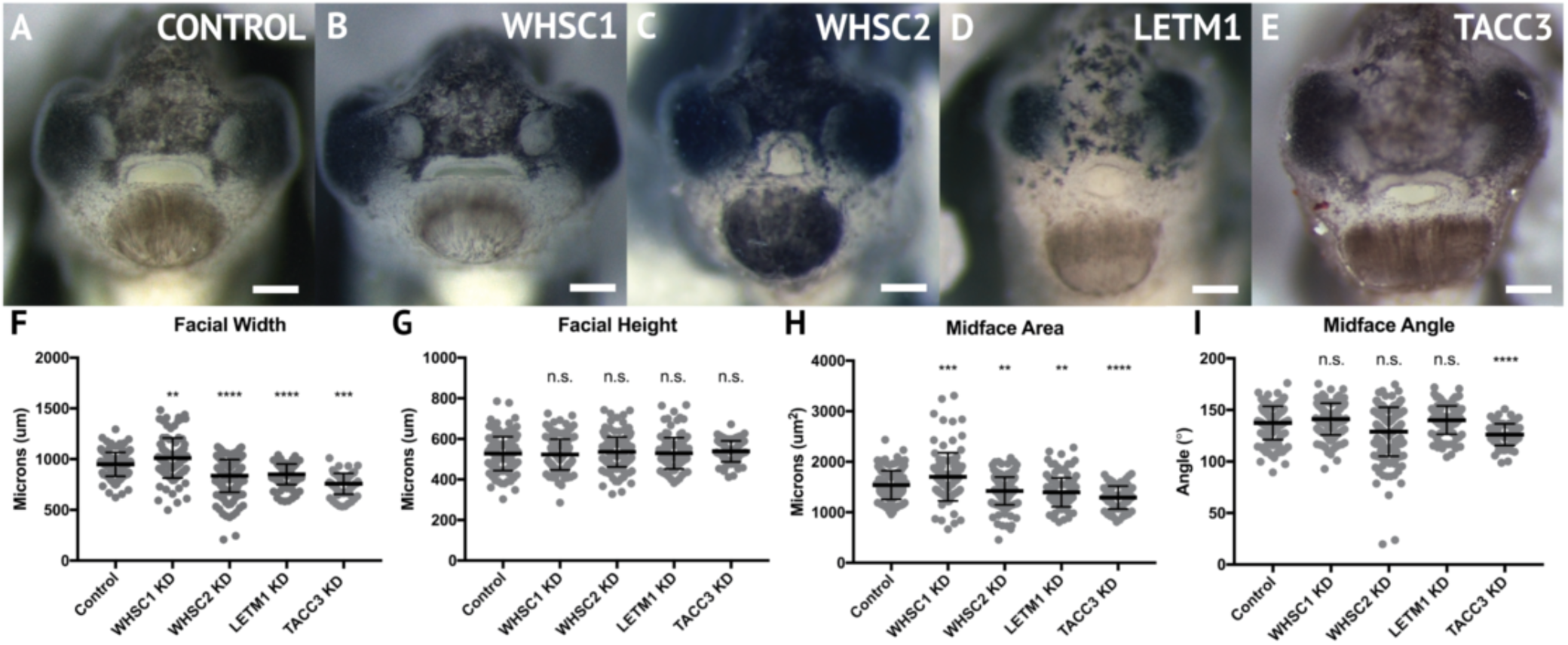
WHS related gene depletion affects craniofacial morphology. (A-E) Frontal views of 3dpf embryos (st. 40) following WHS gene single KD. (F-I) Measurements for facial width, height, midface area, and midface angle. A significant 6.54% increase in facial width and 11.43% increase in midface area were observed for WHSC1 KD. WHSC2 KD caused a 12.01% reduction in facial width and a 6.79% reduction in midface area. LETM1 KD caused a 10.33% decrease in facial width and a 8.49% decrease in midface area. TACC3 KD caused a 21.27% decrease in facial width and a 16.33% decrease in midface area, and an 8.27% decrease in midface angle. Significance determined using a student’s unpaired t-test. (Embryos Quantified: Con MO = 137, WHSC1 MO = 100, WHSC2 MO = 185, LETM1 MO = 115, TACC3 MO = 79.) ****P <0.0001, ***P<0.001, **P<0.01, *P<0.05, n.s., notsignificant. Scalebar = 250μm.

### WHS-affected genes maintain craniofacial cartilage size and scaling

A majority of WHS cases demonstrate defects in cartilage and skeletal formation. Notable examples include underdeveloped ears with reduced or missing cartilage, micrognathia, tooth malformation, short stature, and delayed growth of the head and body [1,19], as well as jaw and throat malformations that significantly impair speech, feeding, and swallowing [1]. The etiology of these co-morbidities is virtually unknown. As craniofacial cartilage and bone are largely derived from cranial neural crest [23], we hypothesized that one or more of these genes may play a critical role in craniofacial cartilage formation. To test this, we performed depletion of WHSC1, WHSC2, LETM1, and TACC3 as described above, in order to survey their impact on scaling and morphology of craniofacial cartilage in *X. laevis* larvae (Fig. 4A-I).

**Fig. 4.**
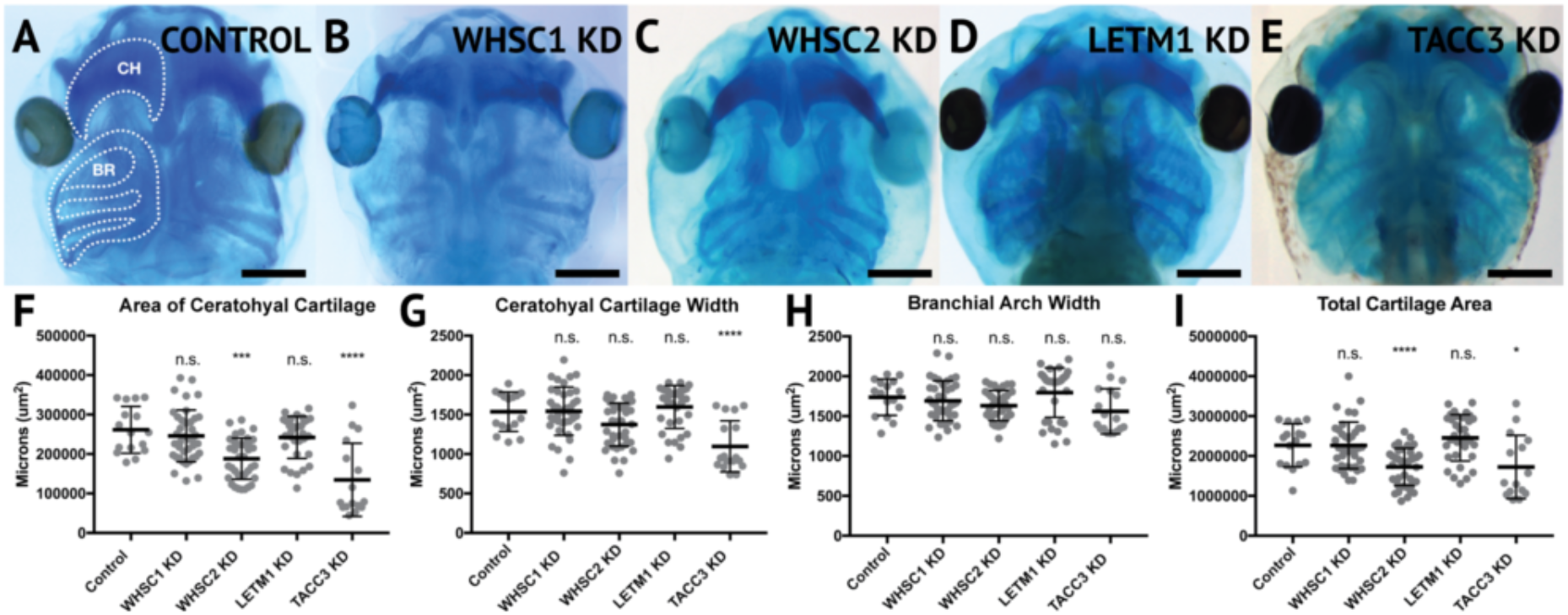
WHSC2 and TACC3 KD impact cartilage morphology. (A-E) Ventral view of 6dpf embryos, stained with Alcian Blue. (F-I) Measurements of area and width of CH, BR width, and total cartilage area. Neither WHSC1 nor LETM1 KD caused a change in measured parameters. WHSC2 KD caused a 27.94% decrease in CH area, and a 23.87% decrease in total cartilage. TACC3 KD caused a 48.5% decrease in area of the CH, a 24.03% decrease in total cartilage, and a 28.58% decrease in CH width. Stat. significance determined by unpaired t-test. (Embryos quant.: Con KD =17, WHSC1 KD = 41, WHSC2 KD = 39, LETM1 KD = 34, TACC3 KD = 11.) ****P <0.0001, ***P <0.001, **P <0.01, *P <0.05, n.s., not significant. Scalebar=250μm.

Depletion of either WHSC2 or TACC3 was sufficient to reduce the combined area of the ceratohyal and branchial arch cartilages (CH and BR, respectively,(4)), in six-day (stage 47) embryos (Fig. 4G). These effects were also explicitly shown in the ceratohyal area alone (Fig. 4F). Ceratohyal cartilage width was also reduced upon TACC3 depletion (Fig. G). Somewhat surprisingly, given the impact of WHSC1 depletion on facial width, its depletion did not increase ceratohyal width or area. Similarly, LETM1 depletion did not reduce cartilage area, despite reduction in overall facial width. These results indicate that WHSC2 and TACC3, genes both within and immediately proximal to the critically-affected locus of WHS, can impact early cartilaginous tissue formation, illustrating a potential avenue through which larger deletions may exacerbate phenotypic severity. Importantly, these effects are demonstrable at 6d post-fertilization, suggesting that early partial depletion of these transcripts produces lingering impacts on craniofacial patterning (first measured at 3d post-fertilization, Fig. 3) that are not ameliorated later in development. We postulated that these persistent craniofacial patterning defects following early depletion of WHS-associated genes may then arise indirectly, from impacts on their embryonic progenitors.

### WHSC1 and TACC3 are critical for normal pharyngeal arch morphology and cranial neural crest cell motility

Given the enrichment of WHS-affected gene transcripts in CNCs post-specification, explicitly in stages that correspond with their migration into the pharyngeal arches (st. 25-35), we hypothesized that their depletion may directly compromise CNC motility. To examine this, we used single-hemisphere injection strategies to generate left-right chimeric embryos, and internally compared pharyngeal arch (PA) migration along control or depleted sides.

Following single-sided WHSC1, WHSC2, LETM1, or TACC3 depletion, embryos were staged to 25-30, fixed, and *in situ* hybridization was again performed against the CNC-enriched transcription factor *Twist*, to visualize migrating PAs(5). Measurements of length, area, and migration were compared to their internal controls. WHSC1 and TACC3 depletion reduced total area of migratory PAs (Figure 5C, H). Further, when WHSC1 levels are reduced, PAs were shorter in length, (Fig. 5D) and their ventral migration distance was reduced compared to paired controls (Fig. 5E). LETM1 and WHSC2 reduction, in contrast, did not result in any significant changes to pharyngeal arch migration *in vivo*. This suggests a role specifically for WHSC1 and TACC3 in maintaining normal migrating pharyngeal arch morphology.

**Fig. 5.**
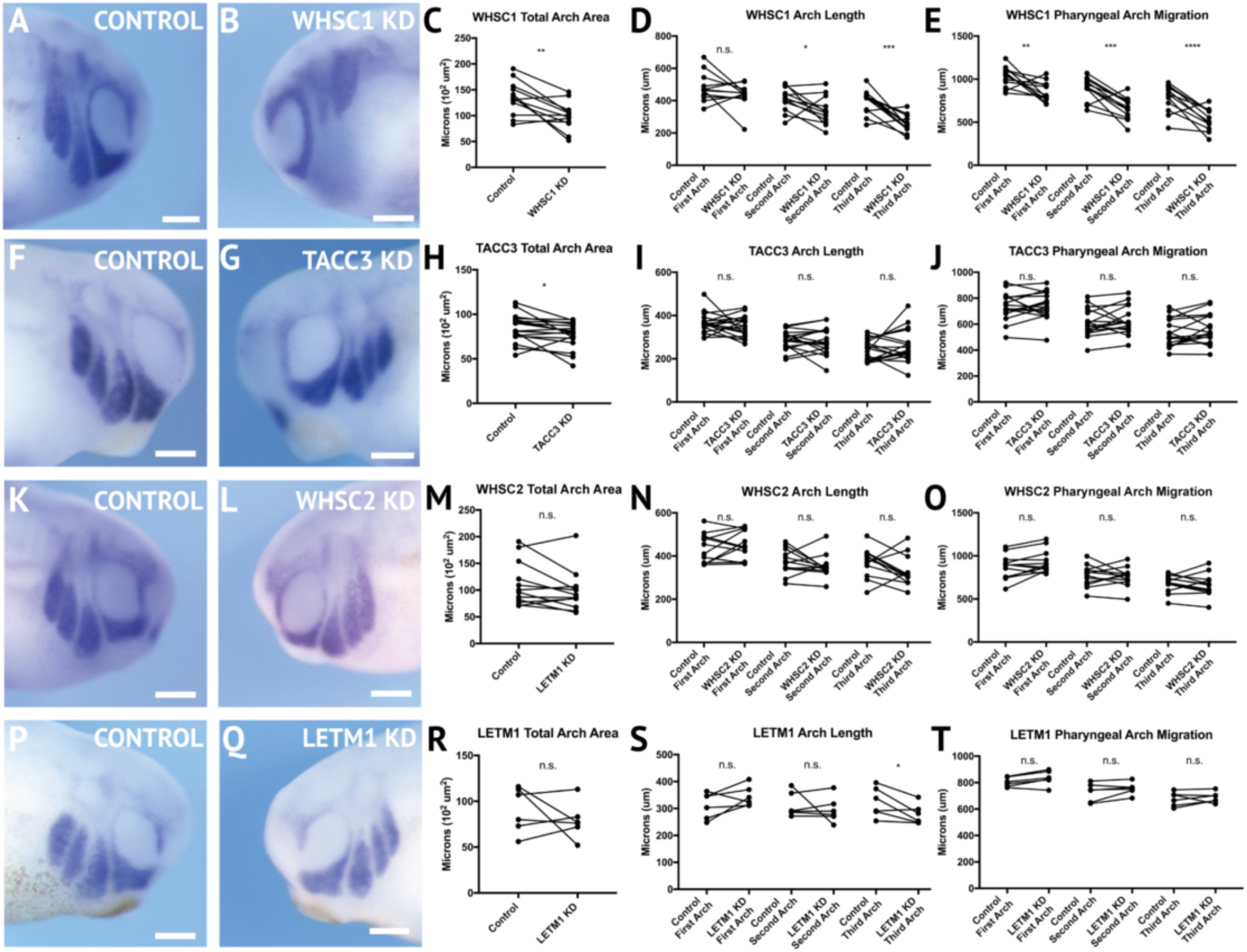
WHSC1 and TACC3 KD decrease CNC migration in *vivo*. (A-B, F-G, K-L, P-Q) Lateral views of embryos (st. 27) following whole mount *in situ* hybridization against TWIST. Each column (A-B, F-G, K-L, P-Q) show lateral views of the same embryo. (C-E, H-J, M-O, R-T) Measurements were taken for total area of the three PA (Arch 1-3 extend anterior to posterior), the length of each arch, and migration distance, as measured from dorsal most tip to the neural tube. (K-T) LETM1 or WHSC2 KD did not significantly affect measured parameters. (F-J) TACC3 KD expression caused an 8.33% decrease in total PA area, but did not affect length or migration. (A-E) WHSC1 KD caused a 23.57% decrease in PA area, and the length of the second and third PAs decreased by 1 4.72% and 31.70%, respectively. The migration distance of the 1st-3rd PAs decreased by 15.75%, 24.04% and 29.29%, respectively. Stat. significance determined by paired t-test. (Embryos quant.: WHSC1 KD = 13, TACC3KD=18, WHSC2KD= 12, LETM1 KD = 19.) ****P <0.0001, ***P <0.001, **P <0.01, *P <0.05, n.s., not significant. Scalebar=250μm.

NCC migration velocity is only one possible contributor to normal PA morphology. Smaller arches, as shown with either WHSC1 or TACC3 depletion, could result from reduced migration rates, or a reduced number of CNCs (indicative of possible separate proliferation defects). To determine whether WHSC1 depletion could specifically impact neural crest migration speed, *in vitro* migration assays were performed as described previously [24,25]. Briefly, whole embryos were injected with either control or WHSC1KD strategies, and their CNCs were dissected prior to their delamination from the neural tube (st. 17). These tissue explants were then cultured on fibronectin-coated coverslips, and trajectories of individual cells that escaped the explant were mapped using automated particle tracking [26,27] to obtain migration speeds(6). WHSC1 depletion resulted in slower individual cell migration speeds compared to controls (Fig. 6B-D, Sup. Video 1). TACC3 also reduced individual neural crest speeds (not shown). We compared these results to those obtained following WHSC2 depletion. As WHSC2 KD was not sufficient to alter PA area or migration in vivo (Fig 5L-O), we hypothesized that cell speed would be unaffected. Interestingly, WHSC2 depletion resulted in a significant increase in speed of CNCs migrating in culture (Fig. 6D). As CNC migration is heavily restricted *in vivo* due to repellent and non-permissive substrate boundaries [28], in addition to the coordinated relationships between neural crest and placodal cell migration [29], it is not surprising that moderate increases in cell velocity *in vitro* would not correspond to an impact on PA migration *in vivo*. In contrast, a deficit in individual cell migration rate, as shown with WHSC1 and TACC3 depletion, would lack comparable compensatory strategies and may more directly delay PA streaming. Thus, we show that WHSC1 depletion alters PA morphology and migration, and that this effect could be directly driven by a reduction in individual CNC migration rates.

**Fig. 6.**
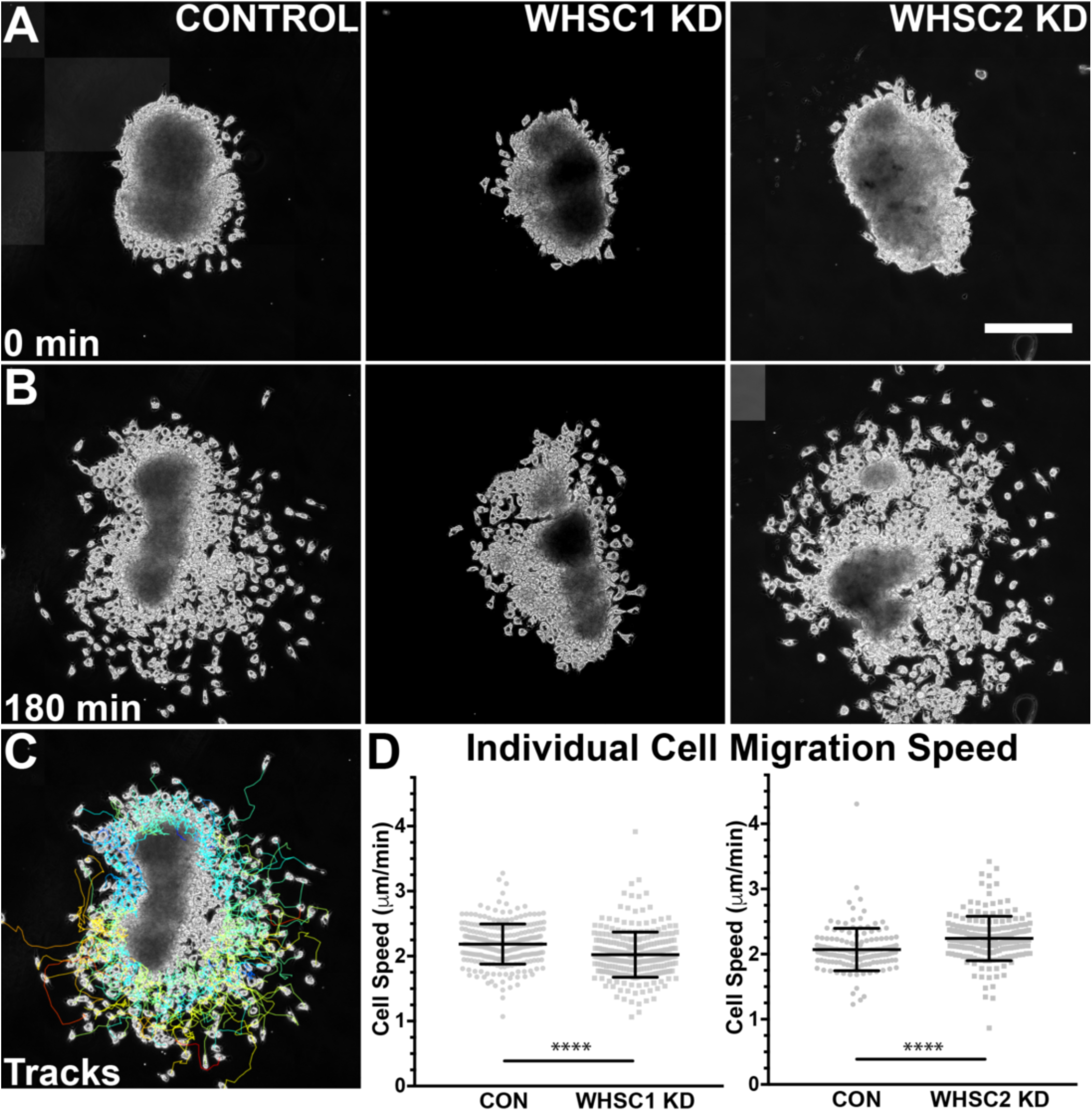
WHSC1 alters CNC migration speeds in vitro. Dissected CNC explants were plated on fibronectin and imaged 3hrs (Phase-contrast, 20x). (A) Representative explants at initial timepoint (0min). (B) Explants after 3hrs migration time. (C) Representative tracks generated by FiJi Trackmate plug-in. (D) Mean track speeds of WHSC1 or WHSC2 KD compared to controls. Stat. signicance determined with unpaired t-test. (Explants quant.: WHSC1 controls = 272 cells, 9explants. WHSC1KD=282cells, 9 explants. WHSC2 controls = 151 cells, 12 explants. WHSC2 KD = 195 cells, 8 explants.) ****P <0.0001. Scalebar = 250μm.

### WHS-related genes impact forebrain morphology

In addition to craniofacial dysmorphism, children with 4p16.3 microdeletions demonstrate mild to profound intellectual disability, with a large majority displaying significant psychomotor and language delays that entirely preclude effective communication [1,19,30]. Larger microdeletions have generally been correlated to more severe intellectual disability and microcephaly, implying that numerous WHS-affected genes may function combinatorially or synergistically to facilitate central nervous system development and cognitive function [19]. Alternatively, this may suggest that genes that are further telomeric within the affected loci could be more impactful contributors to cognitive deficits.

We have largely focused our current efforts to examine the developmental contributions of WHS-affected genes to neural crest migration and craniofacial development, and development of the central nervous system should largely be considered to function distinctly and be examined in future works. However, given the significant craniofacial malformations demonstrated with WHS-associated gene depletion, and the intimate ties between central nervous system and craniofacial development [31,32], we also performed initial characterization of how these WHS-affected genes may singularly contribute to one aspect of neurodevelopment, embryonic forebrain scaling.

To address the impact of WHSC1, WHSC2, LETM1, and TACC3 on forebrain size, we performed half-embryo depletions as above, and examined the outcomes on embryonic brain size. Embryos were injected with single-hemisphere depletion strategies at the 2-cell stage, then allowed to mature to six days (st. 47) prior to fixation. Immunolabeling for alpha-tubulin was carried out to highlight neuronal morphology (7; for experimental workflow, see Fig. S3), and brain areas were compared with paired t-tests between KD and control hemispheres. Forebrain size was significantly reduced with WHSC1, WHSC2, or TACC3 KD (Fig. 7C, F, L). Additionally, control injections did not affect brain size, relative to internal non-injected controls (Fig. S3). WHSC2 depletion caused an additional decrease to midbrain area (Fig. 7F). LETM1 depletion did not impact forebrain sizing (Fig. 7H-I); however, LETM1 deletion is suspected to be the major contributor to seizure development in children with the disorder [20,33]. This only highlights the importance of future characterizations of the cell biological functions of WHS-impacted genes, as it could be expected that LETM1 depletion may instead disrupt normal neuronal excitation, connectivity, or survival [34]. These initial investigations suggest that WHSC1, WHSC2, and TACC3 facilitate normal forebrain development, and perhaps that their depletion is relevant to WHS-associated microcephaly.

**Fig. 7.**
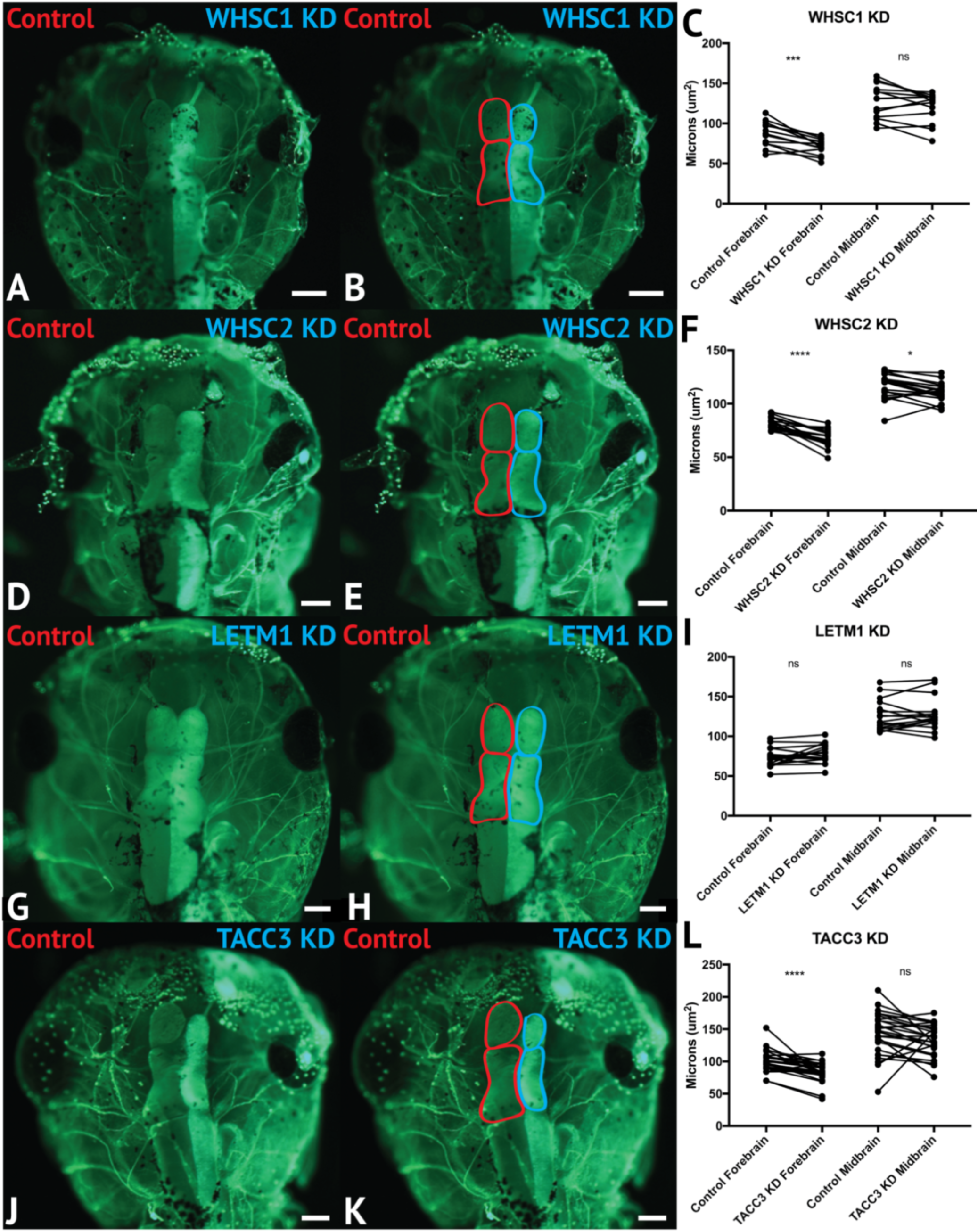
WHSC1, WHSC2 and TACC3 facilitate normal forebrain development. (A-B, D-E, G-H, J-K) Dorsal view of 6 dpf embryos, following half-embryo KDs, with a-tubulin immunolabeling. (B, E, H, K) Dorsal view, with superimposed outlines of forebrain and midbrain structures. Internal control is on left (red), KD is on right (blue). (C, F, I, L) Area of forebrain and midbrain. WHSC1 KD reduced forebrain area by 17.65%. WHSC2 KD reduced forebrain area by 17.33% and midbrain area by 4.14%. LETM1 KD caused no significant change in brain size. TACC3 KD caused a 16.05% decrease in forebrain area. Significance determined using a student’s paired t-test. (Embryos quant.: WHSC1 KD=14,WHSC2 KD = 18,LETM1 KD= 12, TACC3 KD = 26.) ****P <0.0001, ***P <0.001, **P<0.01, *P <0.05,n.s., notsignificant. Scalebar = 250μm.

## Discussion

We have shown that four genes frequently affected in WHS, a human genetic disorder stemming from a heterozygous microdeletion on the short arm of chromosome four, can contribute to normal craniofacial morphogenesis in *Xenopus laevis* (8). We also provide evidence that neural crest migration deficits may significantly contribute to the signature craniofacial dysmorphism of WHS. Specifically, we demonstrate, for the first time, that WHS-associated transcripts are enriched in motile neural crest and contribute to normal craniofacial patterning and cartilage formation (*WHSC2, WHSC1, LETM1*, and *TACC3*). Two of these genes directly impact individual cranial neural crest cell migration (*WHSC1, TACC3*), revealing new basic roles for these genes in embryonic development.

It is increasingly appreciated that full WHS presentation is multigenic [19]; case studies of children with singular gene depletions even in critical regions have historically demonstrated milder syndromic presentations that lack the full range of expected symptoms (intellectual disability, craniofacial abnormalities, seizures, and heart, skeletal, and urogenital defects) [13]. While we have narrowed our examinations to focus on how WHS-affected genes contribute to facial patterning, our findings align well with the idea that WHS presentation is a cumulative product of the impacted locus. While TACC3 and WHSC1 KD impacted all or nearly all examined aspects of craniofacial development at these stages, WHSC1 KD did not produce significant cartilage malformations in isolation, and TACC3 KD narrowed and condensed facial features in a way that appears less analogous to the human ‘Greek Warrior Helmet’ phenotypic presentation.

Of important note, then, WHSC1 depletion was solely able to recapitulate WHS-associated hypertelorism, or facial widening at the level of the eyes and nasal bridge (Figs, 3, 8). As the eyes correspond to the peripheral extrema of the tadpole face, this contributed to a wider face roughly along the axis of the tragion, recapitulating a facial widening demonstrated by 3D morphological mapping of patients with WHS [35]. It is interesting to predict that normal WHSC1 levels may facilitate normal neural crest migration into the face, and in a separate role more explicit to this tissue region, also limit inappropriate proliferation and expansion. In support of this, one of WHSC1’s more established roles is that of an H3K36 methyltransferase, an epigenetic regulator that has been billed as oncogenic, given high levels of dysregulation in some cancer tissues [36,37], and its potential to orchestrate transcriptional programs that drive unchecked proliferation [38]. Other studies report its function to be that of a tumor suppressor, given its high mutation rate in lymphomas [39,40]; additionally, Whsc1 knockout or depletion in zebrafish demonstrated enlarged hearts, brains, and predisposition to swim bladder tumors [41,42], suggesting unchecked expansion of developmental progenitors. As this duality likely partially reflects differential regulation of WHSC1 behavior during development and in the context of oncogenesis, an explicit examination of how WHSC1 functions to regulate tissue expansion and development in the extreme anterior domain may be warranted [43]. Additionally, given that the other three WHS-affected genes instead narrowed facial width and area, this invites further investigation into how these depletions function combinatorially to generate the full signature of WHS craniofacial dysmorphism.

**Fig. 8.**
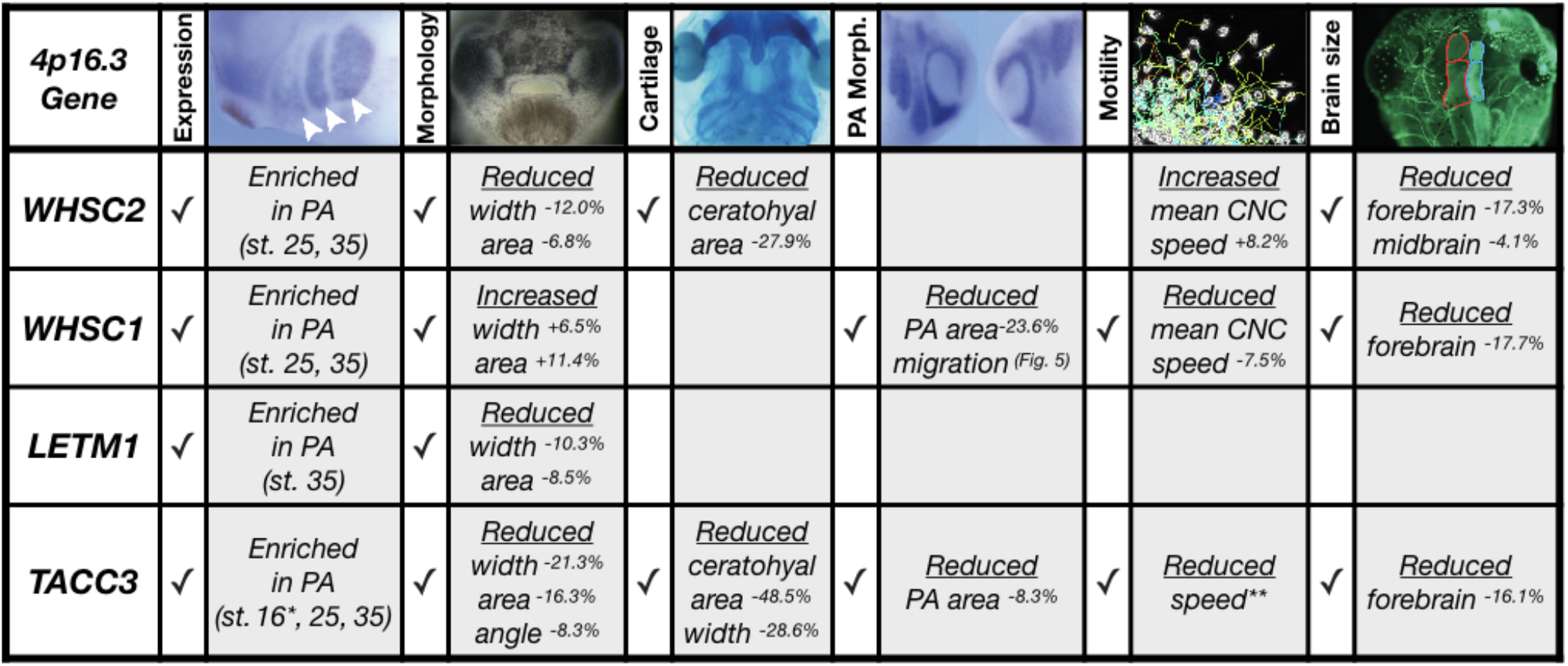
Partial depletion of WHS-affected genes demonstrates numerous impacts on craniofacial development and neural crest migration. Tissues are denoted as affected (checkedbox) if phenotypeswere significantly different from control (p=<.05); see individual figures for data distribution and statistics. (Abbreviations:PA-Pharyngeal Arch) * - Denotes pre-migratory CNC (st. 16) ** - Denotes data in *Bearce e t al., 2018*.

Within that effort, however, it is worthwhile to note that WHS-associated gene depletion in *X. laevis* almost certainly diverges from perfect recapitulation of WHS pathology. *Xenopus* has proven to be an invaluable model for the study of human craniofacial development and disorders [22,44–50], given the highly conserved developmental pathways that drive neural crest migration, differentiation, and craniofacial morphogenesis between systems. Nonetheless, there are gross morphological differences that prevent some direct correlations. It is noteworthy that the CNC that give rise to the ceratohyal cartilage in *Xenopus* will later give rise to far anterior portions of the face, and combine with contributions from the Meckel’s cartilage to form some regions of the jaw [51,52], but equivalent human craniofacial structures undergo distinct development [53]. Loosely, the ceratohyal cartilage in *X. laevis* is formed from CNC of the second PA [44,51]; which in human development will give rise to tissues of the hyoid [53]. Morphological impacts resulting from aberrant development of these tissues, as was shown with either TACC3 or WHSC2 KD (Fig. 4), may then have more direct correlates to human WHS pathology in the context of aberrant pharyngeal development (perhaps leading to speech, feeding and swallowing impairment), rather than explicitly in jaw formation or WHS-associated micrognathia.

Our work has demonstrated consistent enrichment of WHS-associated genes in CNCs, and their necessity for appropriate formation of their derivatives; however, this largely neglects why any of these transcripts may be exceptionally critical in these tissues. This question must be left to some speculation; the precise cell biological roles of all WHS-affected genes warrant much more comprehensive study in the context of embryonic development and cell motility. We have previously summarized some of the known roles of these genes and how they may influence CNC development [54], but a brief summary incorporating recent work is outlined here.

*WHSC2* encodes the gene product Negative Elongation Factor A (NELFA), which functions within the NELF complex to decelerate or pause RNA polymerase II activity [55]. This pausing mechanism is thought to function as a means of synchronizing rapid or constitutive expression of specific transcripts [56–58]. NELF complex components are required during early embryogenesis [59], but their relevance in craniofacial morphogenesis and neural crest migration is entirely unknown. Recent work suggests the NELF complex facilitates cancer cell proliferation and motility, downstream of its regulation of cell-cycle control transcripts [60]. Given that motility and proliferation inherently compete for cytoskeletal machinery [61], the CNC’s somewhat unique need to undergo both rapid expansion and directed motility [62] within the same developmental stages may benefit from these additional levels of coordination, but this remains entirely speculative.

LETM1 localizes to the inner mitochondrial membrane [63], where it acts as a Ca^2+/H+^ anti-porter to regulate Ca^2+^ signaling and homeostasis [33], which can directly affect activity of mitochondrial metabolic enzymes. LETM1 was shown to actively regulate pyruvate dehydrogenase activity, tying its roles directly to glucose oxidation [64]. It’s ubiquitous enrichment across early development (Fig. 2D), and enduring expression within motile CNC (Fig. 2I) might suggest distinct and spatiotemporal metabolic needs during neurulation and craniofacial patterning. Interestingly, NELF complex (containing WHSC2/NELF-A), has been shown to stabilize transcription of fatty acid oxidation-related genes [57], which would suggest dual-depletion of these in areas where they are typically enriched (Fig. 2) may greatly impact metabolic homeostasis. This could be especially damaging in the context of the multipotent CNCs, as metabolism is increasingly demonstrated to perform a commanding roles in determination of cell fate [65–68].

TACC3 is predominantly known as a microtubule regulator. Originally characterized as an essential centrosome adapter during cell division [69,70], its manipulation was more recently shown to impact microtubule plus-end growth in interphase cells and specifically CNCs [71]. It has also demonstrated effects on cytoskeletal mechanics during one form of embryonic cell motility, axon outgrowth and guidance signal response [71,72]. Its significant dysregulation in metastatic cancers [73–75], and roles in mitotic spindle organization [76–79] may allude to additional functions in cytoskeletal coordination of either CNC proliferation or motility, but this remains unexplored. Altogether, it is clear that our current knowledge of how these genes ultimately contribute to embryonic development is sorely lacking, and a basic cell biological examination of WHS-associated gene function within a developmental context is necessary for a better mechanistic understanding of WHS etiology.

Finally, it will also be essential to explore how these genes ultimately synergistically or epistatically regulate WHS pathology. To this aim, our model provides the unique advantage of titratable, rapid, and inexpensive combinatorial depletion of numerous genes, and an intuitive next step will be to perform depletions in tandem that would mirror the genetic perturbations identified from both typical and atypical case studies of WHS. Altogether, our current and ongoing work suggests significant roles for numerous 4p16.3 genes as potent effectors of neural crest-derived tissues and craniofacial morphogenesis.

## Materials and Methods

### *Xenopus* Husbandry

Eggs obtained from female *Xenopus* laevis were fertilized *in vitro*, dejellied and cultured at 13-22°C in 0.1X Marc’s modified Ringer’s (MMR) using standard methods [80]. Embryos received injections of exogenous mRNAs or antisense oligonucleotide strategies at the two or four cell stage, using four total injections performed in 0.1X MMR media containing 5% Ficoll. Embryos were staged according to Nieuwkoop and Faber [81]. All experiments were approved by the Boston College Institutional Animal Care and Use Committee and were performed according to national regulatory standards.

### Immunostaining

5 dpf embryos were fixed in 4% paraformaldehyde in PBS for one hour, rinsed in PBS and gutted to reduce autofluorescence. Embryos were processed for immunoreactivity by incubating in 3% bovine serum albumin, 1% Triton-X 100 in PBS for two hours, then incubated in anti-acetylated tubulin (Sigma, St. Louis MO, USA T7451, 1:500), and goat anti-mouse Alexa Fluor 488 (Invitrogen, 1:1000). Embryos were cleared in 1% Tween-20 in Phosphate Buffered Saline (PBS) and imaged in PBS after removal of the skin dorsal to the brain. Images were taken using a Zeiss AxioCam MRc attached to a Zeiss SteREO Discovery.V8 light microscope. Images were processed in Photoshop (Adobe, San Jose, CA). Area of the forebrain and midbrain were determined from raw images using the polygon area function in ImageJ [82]. Statistical significance was determined using a student’s paired t-test.

### Whole Mount In Situ Hybridization

Embryos were fixed overnight at 4°C in a solution of 4% paraformaldehyde in PBS, gradually dehydrated in ascending concentrations of methanol in PBS, and stored in methanol at −20°C for a minimum of two hours, before *in situ* hybridization, performed as previously described [83]. After brief proteinase K treatment, embryos were bleached under a fluorescent light in 1.8x saline-sodium citrate, 1.5% H2O2, and 5% (vol/vol) formamide for 20 minutes to 1 hour before prehybridization. During hybridization, probe concentration was 0.5 ug/mL.

The *TACC3* construct used for a hybridization probe was subcloned into the pGEM T-easy vector (Promega, Madison, WI). The *Xenopus TWIST* hybridization probe was a kind gift from Dr. Dominique Alfandari (University of Massachusetts at Amherst, MA), subcloned into the pCR 2.1TOPO vector (AddGene, Cambridge, MA). The template for making an antisense probe for *LETM1* was PCR amplified from the reverse transcribed cDNA library, using primer set (5’-CATGGCTTCCGACTCTTGTG, CTAGCTAATACGACTCACTATAGGGCTACAGATGGTACAGAGG-3’), then subcloned into the pCS2 vector (AddGene, Cambridge, MA). Templates for *WHSC1* and *WHSC2* antisense probes were PCR amplified from ORFeomes (European Xenopus Resource Center, UK) with the following primer sets: *WHSC1* forward 5’-CTCATATCCTCGGAAGTCCAGC-3’, *WHSC1* backward 5’-CTAGCTAATACGACTCACTATAGGACCATACAACATCTCCAACAG-3’, *WHSC2* forward 5’-CCTCCGTCATAGACAACGTG-3’, and *WHSC2* backward 5’CTAGCTAATACGACTCACTATAGGAGAGGAGTTGTTGTGTCCAG-3’; these products were cloned into the pDONR223 vector (AddGene, Cambridge, MA). The antisense digoxigenin-labeled hybridization probes were transcribed *in vitro* using the T7 MAXIscript kit. Embryos were imaged using a Zeiss AxioCam MRc attached to a Zeiss SteREO Discovery.V8 light microscope. Images were processed in Photoshop (Adobe, San Jose, CA).

### Depletion and Rescue

Morpholino antisense oligonucleotides (MO) were used to target WHS related genes; injections were performed at the 2-4 cell stage. WHSC2 and TACC3 MOs targeted the translation start site of *Xenopus* laevis *WHSC2* (5-TGTCACTATCCCTCATAGACGCCAT-3) and *TACC3* (5-AGTTGTAGGCTCATTCTAAACAGGA3), respectively. *WHSC1* MO targeted the intron-exon boundary of intron 5 of *Xenopus laevis WHSC1* (5-TGCGTTTTCATGTTTACCAGAGTCT-3) and LETM1 MO targeted the intron-exon boundary of intron 1 of *Xenopus laevis LETM1* (5-ATGACACACAAGTGCTACTTACCCT-3). Standard control MO (5-CCTCTTACCTCAGTTACAATTTATA-3) (Gene Tools, LLC, Philomath OR, USA) was used alongside all conditions.

Knockdown of WHSC2 was assessed by Western blot (Fig. S4). Embryos at stage 35 were lysed in buffer (50 mM Tris pH 7.5, 5% glycerol, 0.2% IGEPAL, 1.5 mM MgCl2, 125 mM NaCl, 25 mM NaF, 1 mM Na3VO4, 1 mM DTT, supplemented with Complete Protease Inhibitor Cocktail with EDTA, Roche). Blotting for WHSC2 was carried out using mouse monoclonal antibody to WHSC2 (Abcam, ab75359, dilution 1:3,000). TACC3 start site MO was validated as previously described [71]. Detection was by chemiluminescence using Amersham ECL Western blot reagent (GE Healthcare BioSciences, Pittsburg PA, USA). The bands were quantified by densitometry using ImageJ [82].

WHSC1 and LETM1 splice site MOs were validated through a Reverse Transcriptase Polymerase Chain Reaction (PCR). Total RNA was extracted by homogenizing embryos (2dpf) in Trizol, and RNA purification was performed according to Qiagen RNA purification protocol. A phenol:chloroform extraction was performed followed by ethanol precipitation. cDNA was synthesized using SuperScript II Reverse Transcriptase. PCR was performed in a Mastercycler using HotStarTaq following the Qiagen PCR protocol. Primers for *LETM1* are as follows; forward 5’-GTACGAGGCTGTGTGCTGAG-3’ and backward 5’-CGGTTTCCACTTCGCTGACG -3’. Primers for *WHSC1* are as follows; forward 5’-GTCGTACAAGAGAAGACGAGTG -3’ and backward 5’-GTCAGTGAAGCAGGAGAAGAAC-3’. Band intensity was measured using densitometry in ImageJ [82] (Fig. S4).

Rescues were performed with exogenous mRNAs co-injected with their corresponding MO strategies. Xenopus ORFs for *WHSC1* and *WHSC2* were purchased from EXRC and gateway-cloned into pCSF107mT-GATEWAY-3’-LAP tag (Addgene plasmid 67618, a generous gift from Todd Stunkenberg). A complete coding sequence of *X. tropicalis LETM1* was purchased from Dharmacon (Lafayette, CO) then subcloned into pCS2+ EGFP vector. Plasmid for *TACC3* cloned into pET30a was a kind gift from the Richter lab (University of Massachusetts Medical School, Worcester, MA), which was subcloned into pCS2. As a start-site MO was utilized to block TACC3 translation, an MO-resistant exogenous mRNA was generated by creating conserved mutations in the first 7 codons. Rescue concentrations are described in Fig. S2.

### Cartilage Staining

At 6dpf, *Xenopus* embryos were anesthetized with benzocaine and fixed in cold 4% paraformaldehyde in PBS overnight. Alcian Blue staining of embryos was performed based on the Harland Lab protocol. Before ethanol dehydration, embryos were bleached under a fluorescent light in 1.8x saline-sodium citrate, 1.5% H2O2, and 5% (vol/vol) formamide for 30 minutes. Embryos were imaged in PBS, using a Zeiss AxioCam MRc attached to a Zeiss SteREO Discovery.V8 light microscope. Images were processed in Photoshop (Adobe, San Jose, CA). Analysis of cartilage structures was performed in ImageJ utilizing the polygon, area, and line functions [82]. Measurements included 1) Total cartilage area measured as the area of the cartilage from the base of the branchial arches, along either side of the cartilage structure, and around the infracostal cartilage. 2) Average ceratohyal cartilage area (outlined cartilage in Fig. 4). 3) Branchial Arch Width was determined by measuring the width of the branchial arch across the widest point. 4) Ceratohyal Cartilage Width was determined using the line function at the widest point on the ceratohyal cartilage. Differences were analyzed by student unpaired t-test.

### Quantifying Craniofacial Shape and Size

Embryos (St. 40, 3d) were fixed in 4% paraformaldehyde in PBS overnight. A razor blade was used to make a cut bisecting the gut to isolate the head. Isolated heads were mounted in small holes in a clay-lined dish containing PBS with Tween. The faces were imaged using a Zeiss AxioCam MRc attached to a Zeiss SteREO Discovery. V8 light microscope. ImageJ [82] software was used to perform craniofacial measurements. These measurements included the: 1) intercanthal distance, which is the distance between the eyes, 2) Face height, or the distance between the top of the eyes and the top of the cement gland at the midline, 3) dorsal mouth angle, which is the angle created by drawing lines from the center of one eye, to the dorsal midline of the mouth, to the center of the other eye, and 4) Midface Area, which is the area measured from the top of the eyes to the cement gland encircling the edges of both eyes. For all facial measurements, Student’s unpaired t-tests were performed between KD embryos and control MO injected embryos to determine statistical relationships. Protocol was lightly adapted from Kennedy and Dickinson (Kennedy and Dickinson, 2014).

### Half-Embryo Injections

Half KDs were performed at the two-cell stage. *X. laevis* embryos were unilaterally injected two times with both WHS gene-specific MO and a GFP mRNA construct. The other blastomere was injected with control MO. Embryos were raised in 0.1X MMR through neurulation, then sorted based on left/right fluorescence. In order to complete pharyngeal arch (PA) visualization, embryos were fixed between stage 21-30 and whole-mount *in situ* hybridization was performed according to the previously described procedure. For brain morphology analysis, embryos were fixed at 6 dpf and prepared for alpha-tubulin immunostaining. Analysis of PAs from *in situ* experiments was performed on lateral images in ImageJ [82]. Measurements were taken to acquire: 1) Arch Area: the area of individual PA determined using the polygon tool. 2) Arch Length: the length of the distance between the top and bottom of each PA. 3) Arch Migration: the ventral most part of the PA to the neural tube. Statistical significance was determined using a student’s paired t-test in Graphpad (Prism).

### Neural crest explants, imaging and analysis

A very helpful and thorough guide to cranial neural crest (CNC) isolation has been described previously [24,25]. We offer only minor modifications here. Embryos (st. 17) were placed in modified DFA solution (53mM NaCl, 11.7 mM Na2CO3, 4.25 mM K Gluc, 2mM MgSO4, 1mM CaCl2, 17.5 mM Bicine, with 50ug/mL Gentamycin Sulfate, pH 8.3), before being stripped of vitelline membranes and imbedded in clay with the anterior dorsal regions exposed. Skin was removed above the CNC using an eyelash knife, and CNCs were excised. Explants were rinsed, and plated on fibronectin-coated coverslips in imaging chambers filled with fresh DFA. Tissues were allowed to adhere forty-five minutes before being moved to the microscope for time-lapse imaging of CNC motility. Microscopy was performed on a Zeiss Axio Observer inverted motorized microscope with a Zeiss 20x N-Achroplan 0.45 NA Phase-contrast lens, using a Zeiss AxioCam camera controlled with Zen software. Images were collected using large tiled acquisitions to capture the entire migratory field. Eight to ten explants, from both control and experimental conditions are imaged at a six-minute interval, for three hours. Explants with epithelial or neuronal contaminant were excluded from analysis. Data was imported to Fiji [27], background subtracted, and cropped to a uniform field size. Migration tracks of individual cells are collected using automated tracking with the Trackmate plug-in [26]. Mean speeds rates are imported to Prism (Graphpad), and compared between conditions using unpaired t-tests. Three independent experiments were performed for each experimental condition.

## ACKNOWLEDGEMENTS

We thank members of the Lowery Lab for helpful discussions, suggestions, and editing. We also thank Eric Snow, Mitchell Lavoie, Katya Van Anderlecht, Katherine Montas, Lucas Ashley, and Molly Connors for technical assistance. We thank Nancy McGilloway and Todd Gaines for excellent *Xenopus* husbandry. We also thank the National Xenopus Resource RRID:SCR-013731 and Xenbase RRID:SCR-003280 for their support.

---

**Supplemental Movie**. Control and WHSC1 KD explants undergo 3hr migration.

**Fig. S1.**
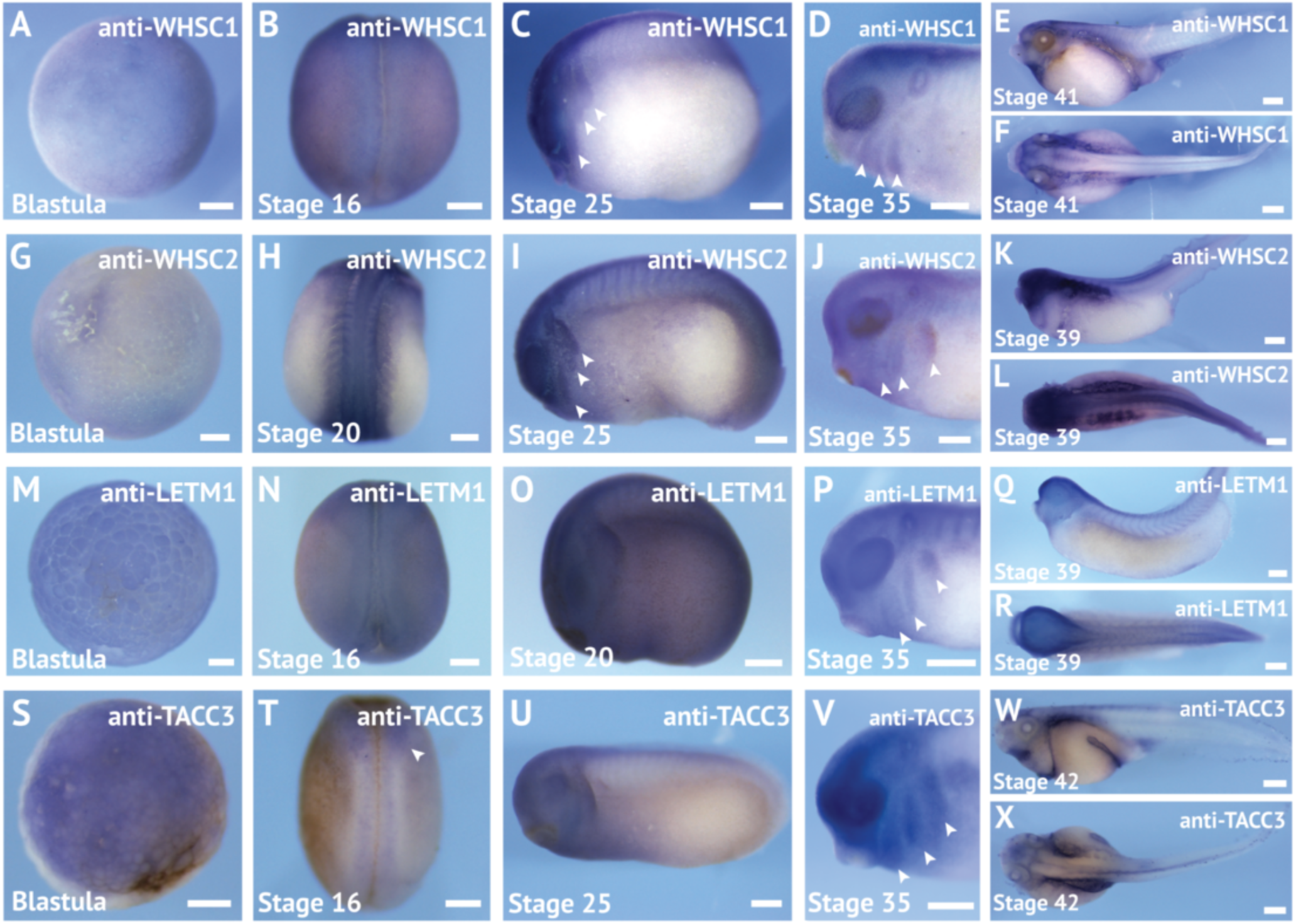
Expression patterns for WHS related genes across early development. *In situ* hybridization utilized (A-F) antisense mRNA probe to WHSC1, (G-L) antisense mRNA probe to WHSC2, (M-R)antisense mRNA probe to LETM1, and (S-X)antisense mRNA probe to TACC3. Embryos shown at blastula stage (A,G,M,S),in dorsal view at stage 16-20 (B,H,N,T), in lateral view at stage 20-25 (C,I,O,U), detail of lateral anterior region at stage 35 (D,J,P,V), and in both lateral and dorsal views from stages 39-42 (E,K,Q,W and F,L,R,X). Scalebar is 250μm.

**Fig. S2.**
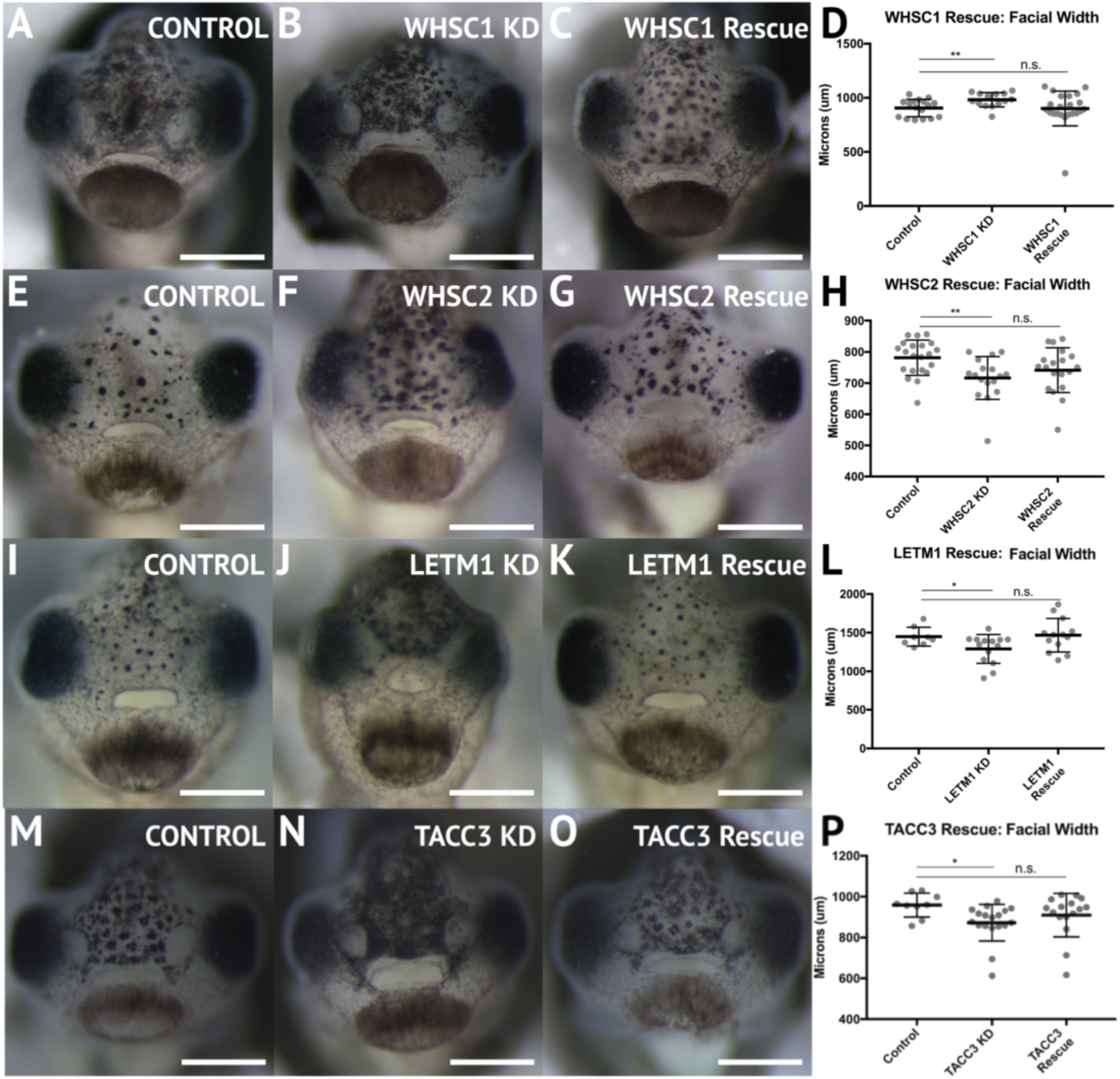
Craniofacial defects caused by WHS-associated gene KD are rescued by co-injection of exogenous mRNA co-expression. Facial widths from control, depletion, or rescue strategies was measured in tadpoles (st. 40). Row 1: Embryos injected with A) control MO (n=17), B) 10ng WHSC1 MO (n=14), or C) 10ng of WHSC1 MO and 250 pg of WHSC1 exogenous mRNA. D) Comparisons of facial width showed an 8.76% increase in facial width with WHSC1 KD, which was rescued by WHSC1 mRNA co-injection. Row 2: Embryos injected with E) control MO (n=21), F) 10ng WHSC2 MO (n=17), or G) both 10ng of WHSC2 MO and 250 pg of WHSC2 mRNA (n=19). H) WHSC2 knockdown caused an 8.37% reduction in facial width, which was rescued by exogenous WHSC2 mRNA co-injection. Row 3: Embryos injected with I) control MO (n=10), J) 20ng of LETM1 MO, or K) 20ng LETM1 MO and 1500pg of LETM1 mRNA (n=11). L) KD of LETM1 caused a 14.95% decrease in facial width, and was rescued by co-injection of exogenous LETM1 mRNA. Row 4: Embryos injected with M) control MO (n=9), N) 20ng of TACC3 MO (n=18), or O) 20ng of TACC3 morpholino and 1000pg of TACC3 mRNA (n=16). P) TACC3 KD resultedina9.01%decrease in facial width,and was rescued by TACC3 mRNA co-injection. Significance determined using a student’s unpaired t test. ****P <0.0001, ***P <0.001, **P <0.01, *P <0.05, n.s., not significant. Scalebar is 250μm.

**Fig. S3.**
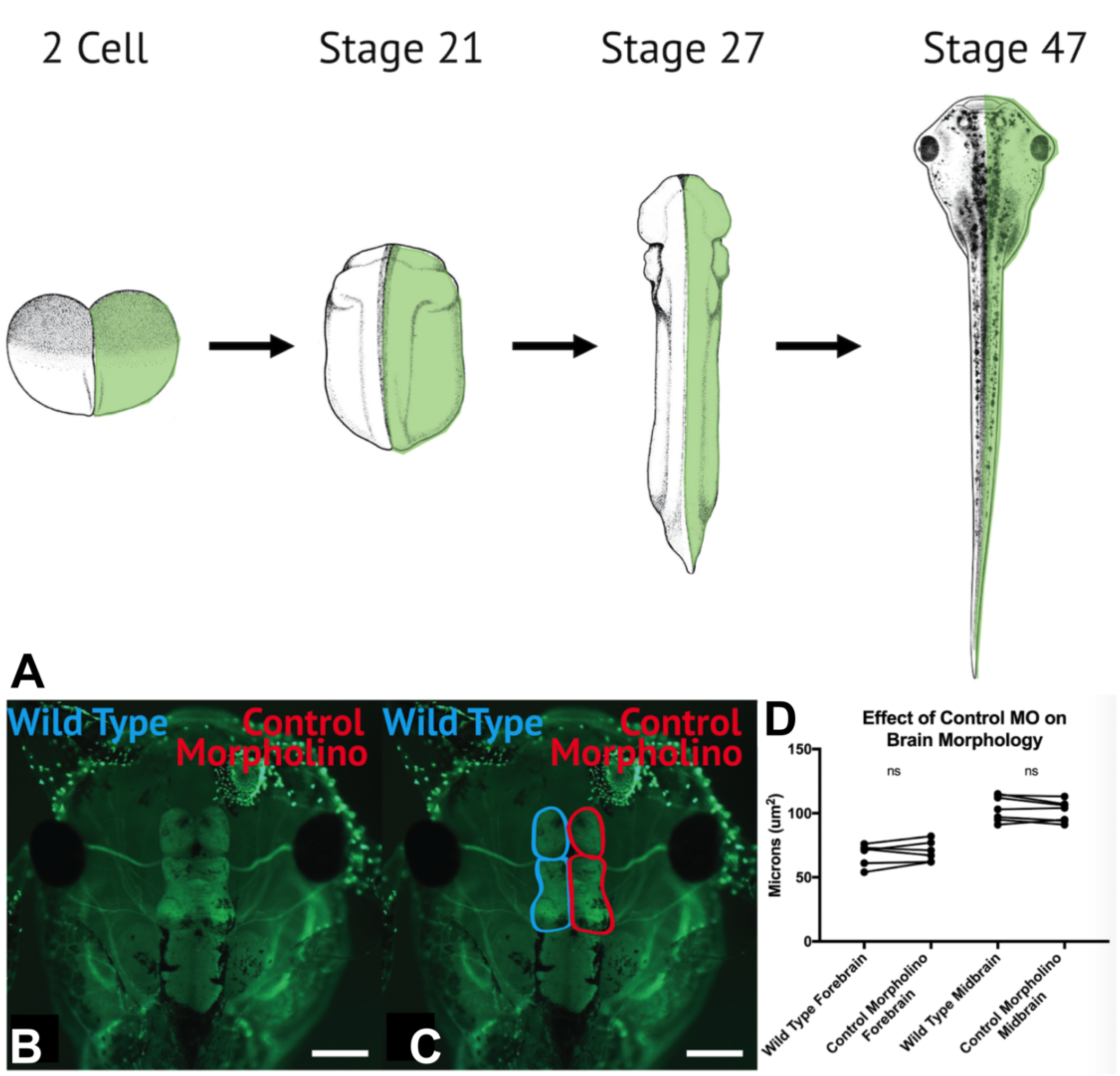
Half embryo knockdown can be utilized for analysis of brain morphology and neural crest cell migration in vivo. (A) At the 2-cell stage, a single blastomere is injected with WHS-associated gene MOs and exogenous eGFP mRNA.After neurulation, embryos are sorted based on left or right eGFP fluorescence, to determine side of depletion. Embryos were raised to st. 47, then fixed and stained with a-tubulin to highlight neuronal morphology. (B-D) Control MO does not significantly impact brain size, compared to non-injected hemispheres (a paired internal control).

**Fig. S4.**
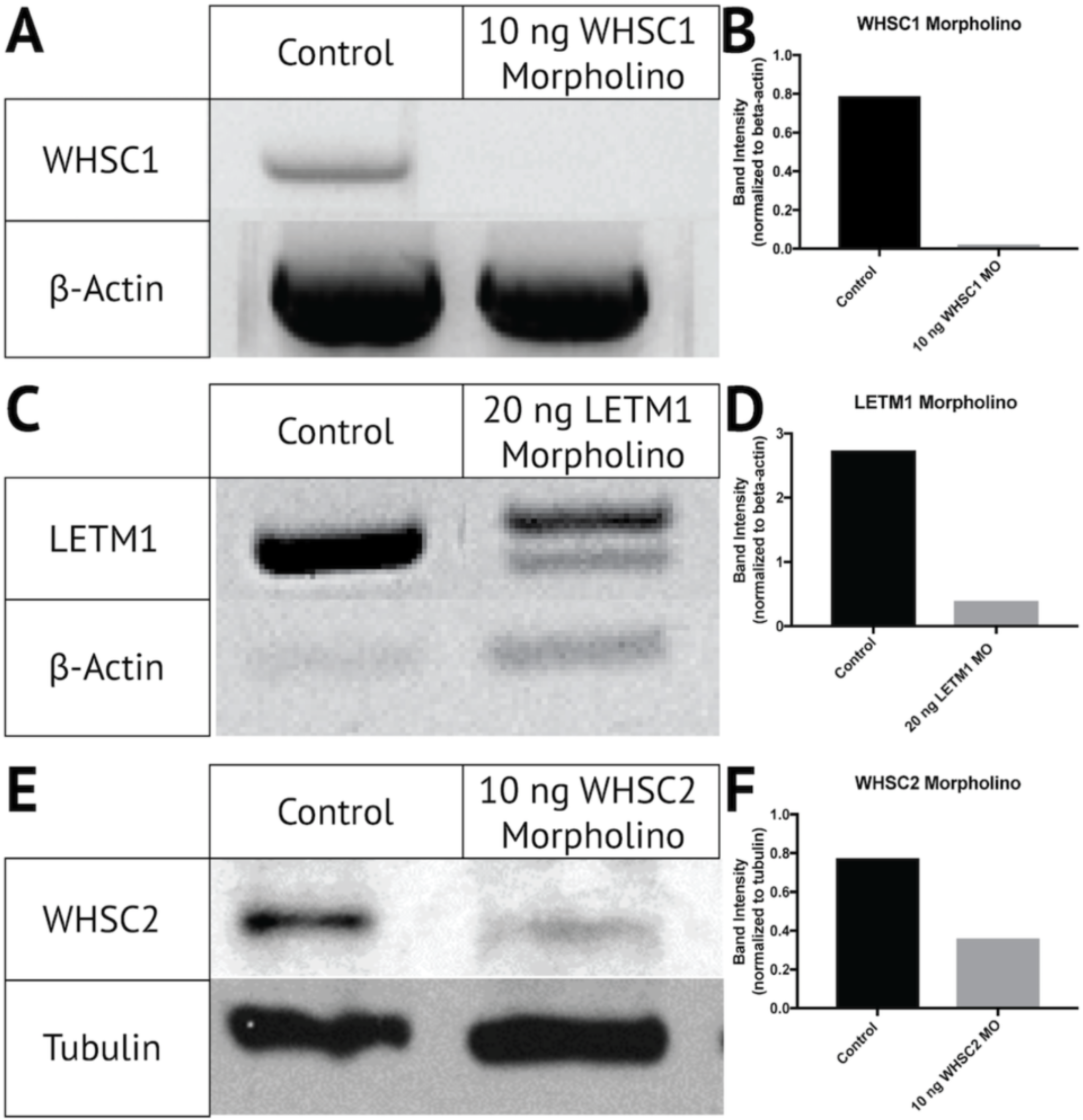
Validation of WHS MOs. A-B) Gel of polymerase chain reaction (PCR) that shows injection of 10ng of WHSC1 MO causes a greater than 90% reduction in WHSC1 mRNA at 2dpf. C-D) Gel of PCR showing injection of 20ng of a MO targeted against LETM1 causes an 80% decrease in LETM1 mRNA 2dpf. Note two bands appear, providing confirmation of splice site error and size shift. E-F) Western blot showing 10ng injection of a MO targeted against WHSC2 results in a greater than 50% reduction in WHSC2 protein by 2dpf. Bar graphs (B,D,F) depict densitometry of gels (B,D) or blot(F)shown, but is consistent across triplicate results.

